# A novel nitric oxide (NO)-dependent ‘molecular switch’ mediates LTP in the *Octopus vulgaris* brain through persistent activation of nitric oxide synthase (NOS)

**DOI:** 10.1101/2024.12.25.630295

**Authors:** Ana Luiza Turchetti-Maia, Tal Shomrat, Flavie Bidel, Naama Stern-Mentch, Nir Nesher, Binyamin Hochner

## Abstract

Cephalopods are a renowned example of the independent evolution of complex behavior in invertebrates. The octopus’s outstanding learning capability is a prominent feature that depends on the vertical lobe (VL). Previously, we found that the synaptic input into the VL exhibits robust activity-dependent long-term potentiation (LTP) mediated by molecular processes that are only partially understood. Here, we reveal that the VL LTP is mediated by nitric oxide (NO). In contrast to the prevailing dogma, in the octopus VL, NO does not mediate LTP *induction*, as tetanization-induced LTP occurs even in the presence of NO-synthase (NOS) inhibitors. Remarkably, however, NOS inhibitors block the long-term presynaptic expression of LTP, and high doses of NO donor induce short-term synaptic potentiation, suggesting that a persistent elevation of NO concentration mediates LTP *expression*. Moreover, in a distinct group of synapses, NOS inhibitors also disrupted LTP maintenance, as following drug washout, a high-frequency stimulation reinstated full LTP, suggesting that NO is also involved in maintaining LTP. We propose a novel molecular-switch mechanism whereby a positive feedback loop of NO-dependent NOS reactivation mediates persistent NOS activation, thus providing an LTP *maintenance* mechanism. Subsequently, retrograde NO diffusion facilitates presynaptic transmitter release, driving LTP *expression*. These findings demonstrate how evolutionary adaptation of molluscan molecular mechanisms has contributed to the emergence of the advanced cognitive abilities observed in octopuses.

## Introduction

Octopuses and other modern coleoid cephalopods (squid and cuttlefish) are outstanding examples for the independent evolution of complex motor (*Levy and Hochner 2017; Levy et al. 2017; Nesher et al. 2020; Hochner and Zullo et al. 2023*) and cognitive behaviors in invertebrates (*Hanlon and Messenger 2018; Turchetti-Maia et al. 2017; Shomrat et al. 2015; Wells 1978; Amodio et al. 2019*). As such, they are renowned for possessing the largest nervous system of all invertebrates, consisting of approximately half a billion neurons (*Young 1971*). The vertical lobe (VL), an area located in their central brains that controls learning and memory (*Figure 1A and B*), matches in its gross morphological and neurophysiological characteristics the connectivity scheme of feedforward fan-out fan-in association/classification networks (*Shomrat et al. 2011; Babadi and Sompolinsky 2014; Aso et al. 2014; Litwin-Kumar et al. 2017*). In the octopus, the fan-out input layer consists of axons projecting out of 1.8 million superior frontal lobe neurons (SFLn) (*Figure 1B and C*); these axons innervate *en passant* 25 million small amacrine interneurons (AM). A recent connectome study (*Bidel and Meirovitch et al. 2023*) found that in contrast to previous views (*Gray 1970; Young 1971*), the AMs are not a single homogeneous group of interneurons (see connectivity scheme, *Figure 1C*). The large majority (89.3% of the 25 million VL neurons) were termed “simple” AMs (SAMs) as they project a single non-bifurcating neurite into the neuropil. Surprisingly, each SAM neurite receives only a single synaptic input through a large SFLn axonal varicosity as it crosses the SFL tract. A very small group of newly discovered AM interneurons (∼1.6%) are classified as complex AMs (CAMs) due to their bifurcating neuritic trees, which receives multiple synaptic inputs from numerous SFLn axons as well as from multiple SAMs. The SAMs and CAMs then converge onto the dendritic processes of the VL output layer, comprising 65,000 large efferent neurons (LN) (*Figure 1C*). These connectome findings, along with a recent comprehensive histolabeling study (*Stern-Mentch et al. 2022*), led Bidel and Meirovitch et al. (*2023*) to propose that the VL circuitry is organized in two parallel and interconnected feedforward networks: an excitatory network emerging from the cholinergic SAMs and an inhibitory network likely emerging from the GABAergic CAMs (*Figure 1C*). Indeed, inhibitory postsynaptic potentials (IPSPs), in addition to the cholinergic excitatory EPSPs, were recorded in the LNs (*Shomrat et al. 2011; Bidel and Meirovitch et al. 2023*).

**Figure 1.**
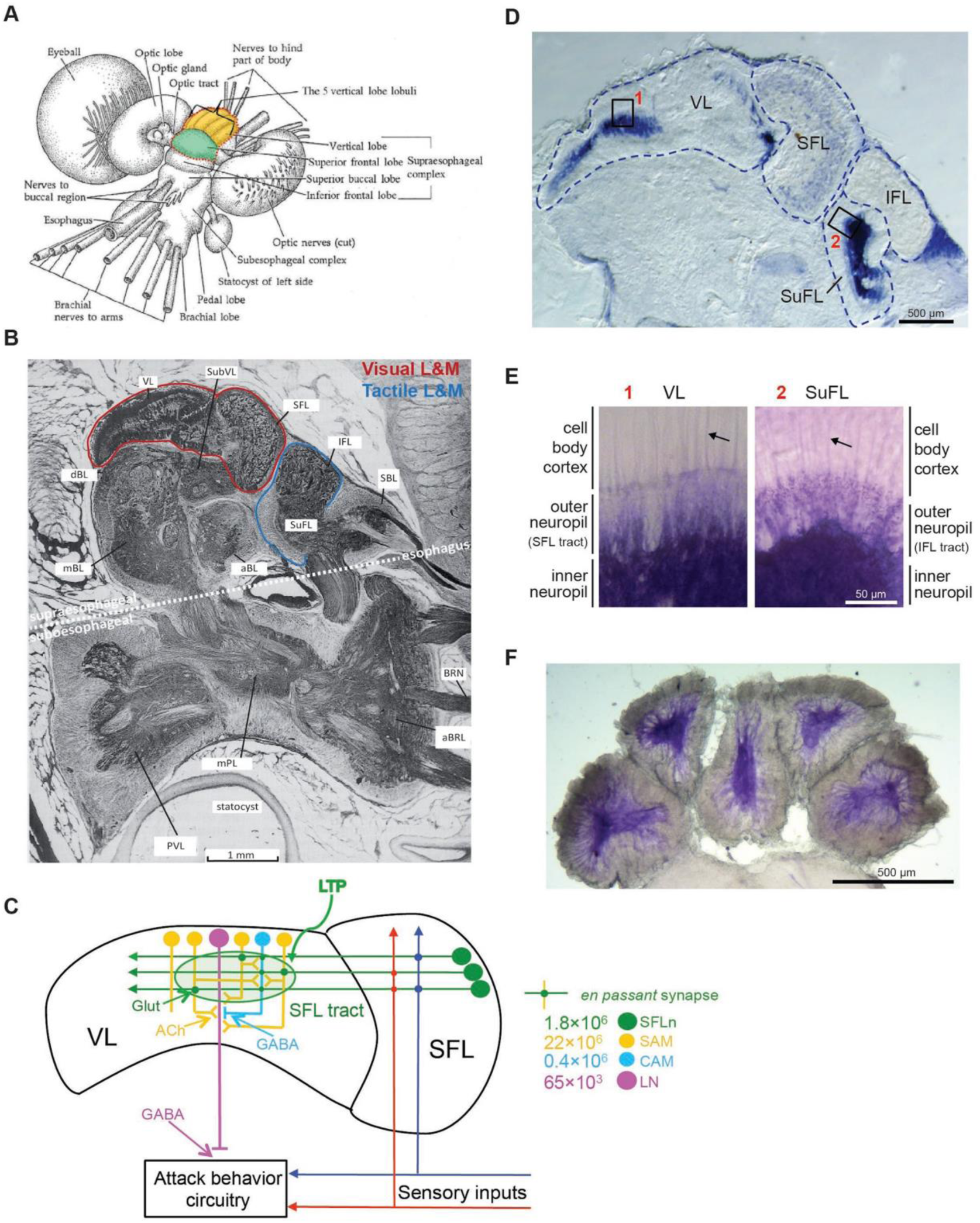
The areas in the octopus brain that are involved in learning and memory are localized in district lobes which express intense NOS activity. (A) The centralized brain of *Octopus vulgaris* in anterior-dorsal view. The supraesophageal brain complex includes the superior frontal lobes (SFL, green) and the vertical lobe (VL, orange), which consists of five cylindrical lobuli (modified from Brusca and Brusca, 1990). (B) A sagittal section of the sub- and supraesophageal brain complexes (modified from Nixon and Young, 2003) showing the location of the SFL-VL system (visual L&M system; red line) and the IFL-SubFL system (chemo-tactile L&M system; blue line) (VL = vertical lobe; SubVL = subvertical lobe; SFL = superior frontal lobe; IFL = inferior frontal lobe; SubFL = subfrontal lobe; SBL = superior basal lobe; aBL = anterior basal lobe; mBL = medial basal lobe; dBL = dorsal basal lobe; BRN = brachial nerve; mPL = medial pedal lobe; PVL = palliovisceral lobe. (C) A schematic wiring diagram of the neural elements of the VL system, showing the basic connectivity (based on Bidel and Meirovitch et al. 2023) (SFLn = SFL neurons; SAM = simple amacrine; CAM = complex amacrine; LN = large efferent neuron). The type of transmitters, the synaptic sites, the number of cells and the site of LTP are indicated by the corresponding colors. (D-F) Histochemical detection of NO-synthase (NOS) activity by NADPH diaphorase staining of supraesophageal brain mass slices (modified from Stern-Mentch et al. 2022). (D) Sagittal section; note intense labeling of the neuropil in VL (1) and SubFL (2). (E) Higher magnification of the respective areas in D. Arrows point to labeling of the AMs’ primary neuritic trunks. (F) Transversal section through the five lobuli composing the VL, showing intense labeling of their neuropil.

As universally found in learning and memory networks, in the octopus VL there is an extremely robust activity-dependent long-term potentiation (LTP) at the SFL-to-AM glutamatergic synapses (*Hochner et al. 2003; Shomrat et al. 2011*). This LTP was proven to be important for learning as “saturating” the VL LTP by global tetanization interfered with the acquisition of long-term memory (*Shomrat et al. 2008*). The octopus activity-dependent LTP was found to be NMDA-independent and its expression involves facilitation of transmitter release from the presynaptic sites of the SFL-to-AM glutamatergic synapses (*Hochner et al. 2003; Shomrat et al. 2011*). Interestingly, these properties are similar to those of the mossy fiber-to-pyramidal cell synapses in the hippocampal CA3 region (*Nicoll and Schmitz 2005*). Approximately half of the synaptic connections in the VL exhibit a key property required for Hebbian plasticity, as LTP induction was blocked when postsynaptic responses were entirely inhibited by kainate-like receptor antagonists (*Hochner et al. 2003*). The postsynaptic-dependent induction combined with the presynaptic LTP expression, led to the hypothesis that a retrograde messenger, such as nitric oxide (NO), which can easily diffuse from the postsynaptic to the presynaptic side of the synapse, plays a role in this form of LTP (*Regehr et al. 2009; Hardingham et al. 2013*).

In the current study, we explored the molecular processes that underlie *induction*, *expression*, and *maintenance* of LTP in the octopus VL. In addition to the mechanistic insights discussed above, previous studies have suggested that NO is involved in visual and tactile learning in *Octopus vulgaris* (*Robertson et al. 1995; Robertson et al. 1996; Robertson et al. 1994*). Furthermore, NO is a well-established mediator of synaptic plasticity in mollusks, particularly in processes associated with more complex forms of learning (*Briskin-Luchinsky et al. 2018; Katzoff et al. 2002; Kemenes et al. 2002; Korshunova and Balaban 2014; Susswein et al. 2004*) than that of the simpler form of learning like in the defensive reflex of the *Aplysia* (*Kandel et al. 2014*).

We found that, contrary to the prevailing dogma, NOS inhibitors did not block LTP *induction*, but, surprisingly, inhibited LTP *expression* and, in some synaptic connections, particularly within the medial lobule, also disrupted LTP *maintenance*. This unprecedented mechanism involves a profound adaptation of the molluscan NO neuromodulation system, which led to the selection of a ‘molecular memory switch’ that in principle resembles the kinase autophosphorylation mechanism proposed by Lisman (1985). We suggest that in the octopus VL, NO-dependent NOS reactivation establishes a positive feedback loop that sustains persistent NOS activation, underpinning LTP *maintenance*. In parallel, NO retrogradely induces a NO-dependent facilitation of presynaptic transmitter release, driving LTP *expression*.

## Results

### The neuropil of the vertical and sub-frontal lobes are intensely labeled for NOS activity

In a previous study, we tested for the presence of NOS activity in the supraesophageal part of the brain employing histochemical techniques (*Stern-Mentch et al. 2022*). Briefly, using the NADPH-diaphorase method (*Hope et al. 1991; Moroz 2000*) we discovered a particularly intense labeling in the neuropils of the VL and the subfrontal lobe (SuFL). These areas are associated with visual and tactile learning, respectively (*Figure 1D-F*), and also share a similar anatomical organization (*Young 1971*). Dense labeling was found in the inner neuropil zones of the five lobuli of the VL (*Figure 1F*), where the synaptic connections between the SAMs, CAMs and the LNs reside (*Gray 1970; Bidel and Meirovitch et al. 2023*). The outer neuropil (*Figure 1E*) was more sparsely stained because, in this region, unlabeled axons that run in the SFL tract make sparse *en passant* connections with SAMs and CAMs neurites (*Gray 1970; Bidel and Meirovitch et al. 2023*). The non-bifurcating neurites of monopolar SAM interneurons are likely those that can be seen crossing the tract in faintly stained AMs trunks (arrows; *Figure 1E*), suggesting the presence of NOS definitely in the SAMs and possibly other VL neurons (*Bidel and Meirovitch et al. 2023*) (see Discussion).

### NO does not mediate LTP *induction*

Physiological experiments on the VL slice preparations were used to test whether NOS inhibitors affect the three elemental stages of LTP: *induction, expression*, and *maintenance*. Accordingly, to test NO involvement in *induction,* NOS inhibitors were administered before and during LTP induction by a high-frequency (HF) stimulation (see details in *Figure 2—figure Supplement 1* in Supplemental methods and analysis).

The results in Figure 2A indicate that NO is not involved in the induction process, as in the presence of the two specific NOS inhibitors, L-NAME (10 mM) or L-NNA (10 mM), HF stimulation induced lower levels of potentiation relative to that obtained in the presence of D-NAME (10 mM) (the inactive enantiomer of L-NAME). However, in contrast to D-NAME, washing out the two inhibitors was clearly followed by a monotonical increase in the magnitude of the postsynaptic field potential normalized to the tract potential amplitude (note that usually we show the second fPSP of the paired pulse test stimulation. i.e., fPSP2/TP2; *Figure 2—figure Supplement 1B*). Importantly, a second HF stimulation (at 100 min) did not lead to facilitation, suggesting LTP saturation. These results indicate that LTP was induced in the presence of the specific NOS inhibitors, but yet, LTP expression was curtailed by the inhibitors and was gradually exposed during the washout of the inhibitors. This supports the premise that NOS is involved in either the expression and/or the development of LTP maintenance mechanisms. As shown in Figure 2B, L-NAME and D-NAME, but not L-NNA, reversibly affect the amplitude of the tract potential (TP) and therefore suggests that this effect is not NO specific. Note, however, that this effect of D- and L-NAME contribute to the reduction in fPSPs amplitude (see below).

**Figure 2.**
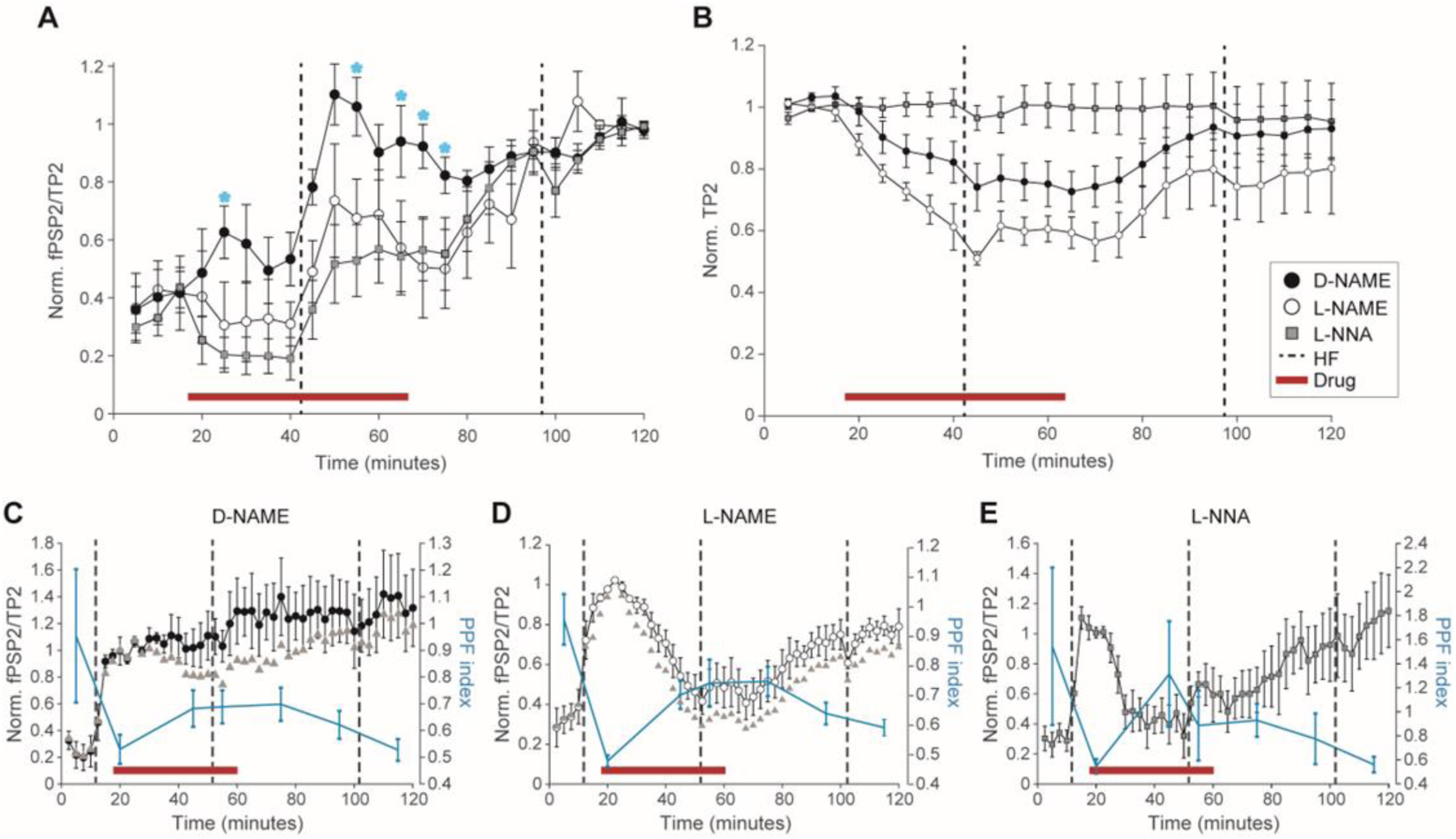
NOS inhibitors do not block LTP *induction* but do block LTP *expression*. (A) Exposing slice preparations to NOS inhibitors (red bar) (L-NNA 10 mM, n=7; L-NAME 10 mM, n=5), but not to the inactive enantiomer control (D-NAME 10 mM, n=5), interfere with the full onset of LTP following HF stimulation (vertical dashed line) in the presence of the drugs. The potentiation was significantly lower with the inhibitors (blue asterisks indicate P < 0.05 between L-NAME and D-NAME). Note the slow recovery of the fPSP following washout of the inhibitors, and the lack of further potentiation following the second HF, suggesting that induction occurred in the presence of NOS inhibitors. (B) The effect of the same three drugs on the TP amplitude. L-NNA, in contrast to both D- and L-NAME, did not affect the presynaptic action potentials. (C-E) The effects of the drugs on LTP expression. (C) D-NAME (10 mM, n=6), (D) L-NAME (10 mM, n=5) and (E) L-NNA (10 mM, n=7). Each panel show the effect of the drugs on fPSP2/TP2 and PPF index (blue; right vertical axes). The PPF index estimates the effect of the drugs on paired pulse facilitation (PPF index = (fPSP2/TP2-fPSP1/TP1)/fPSP2/TP2 (see *Figure 2—figure supplement 1*). The fPSP2/TP2 amplitude is normalized to that of the LTP levels before the administration of the drugs. The drugs perfusion started 10 min after LTP induction (red bar). The gray triangles in C and D depict the “true” effect of the drugs on fPSP by correcting fPSP2/TP2 relative to the reduction of TP2 amplitudes (see B). Note that L-NAME and L-NNA blocked LTP expression in parallel with a reversible increase in the PPF index (see text). D-NAME (C), also reversibly increased the PPF index suggesting an inhibitory effect of D-NAME which is exposed when corrected for the D-NAME effect on the TPs amplitude (gray triangles). A second HF was given 7.5 min before starting drugs washout. A third HF was given toward the end of the recovery phase to check for residual LTP. Data shown as mean ± SEM, dashed vertical lines mark HF stimulation. The average and SEM of the PPF index were calculated over four time points (10 min) around the displayed points.

These results led us to postulate that in contrast to the prevailing dogma, implicating NO in LTP *induction* (*Regehr et al. 2009; Hardingham et al. 2013*), in the octopus VL, NO is involved in LTP *expression* and/or *maintenance*.

### Persistent NOS activation mediates the long-term *expression* of the VL LTP

To test the involvement of NO in LTP *expression* we added the NOS inhibitors after LTP induction (*Figure 2C-E*). L-NAME (10 mM) and L-NNA (10 mM) inhibited the facilitated fPSP when added 10 min after HF stimulation (*Figure 2D,E*). The inactive enantiomer D-NAME (10 mM) did not cause an apparent inhibition (*Figure 2C*). A second HF stimulation, before rinsing out the drugs, did not overcome the inhibition (*Figure 2 D,E*). The third HF stimulation, after drug washout, did not induce additional LTP, thus supporting a recovery of LTP expression.

The average PPF-index (see Methods, and *Figure 2*—*figure Supplement 1*) during the various stages of the experiments (*Figure 2C-E* in blue, and the right vertical axis) show that inhibition was accompanied by a significant increase in the PPF-index, suggesting that the inhibition of LTP *expression* involves a reversible reduction in the probability of transmitter release. Although D-NAME apparently did not inhibit fPSP2/TP2, it did cause a reversible increase in the PPF-index (*Figure 2C*). This effect might indicate that D-NAME has a weak inhibitory effect, especially at the high concentrations we used. Indeed, when we compensated for the effect of D-NAME on the TPs amplitude (see *Figure 2B*) and assuming that the linear relationship between TP and fPSP amplitudes is maintained (*Shomrat et al. 2011*), a weak inhibition by D-NAME is revealed (*Figure 2C*; triangles). As the specific inhibitory effect of L-NAME on fPSP is much stronger than on TPs amplitude the correction showed only a small increase in the level of the apparent inhibition (*Figure 2D*; triangles). The weak D-NAME inhibition relative to that of L-NAME is expected as the drugs are designed to compete on NOS binding site of L-arginine (*Víteček et al. 2012; Griffith and Stuehr 1995*), and thus further support the involvement of NOS in the VL LTP *expression,* irrespective of their similar nonspecific effect on TPs. As L-NNA seems to be more NOS specific, with no apparent effect on TPs amplitude, we used this drug in the remaining of the experiments.

We next investigated if NO is similarly involved in the later phases of LTP, by exposing the VL slices to a NOS inhibitor at different time intervals after LTP induction (*Figure 3A-C*). L-NNA (10 mM for 45 min) was able to block LTP expression when tested at least 4 h after the LTP induction (*Figure 3C*), suggesting that persistent activation of NOS is the mechanism mediating the long-term *expression* of the VL LTP.

**Figure 3.**
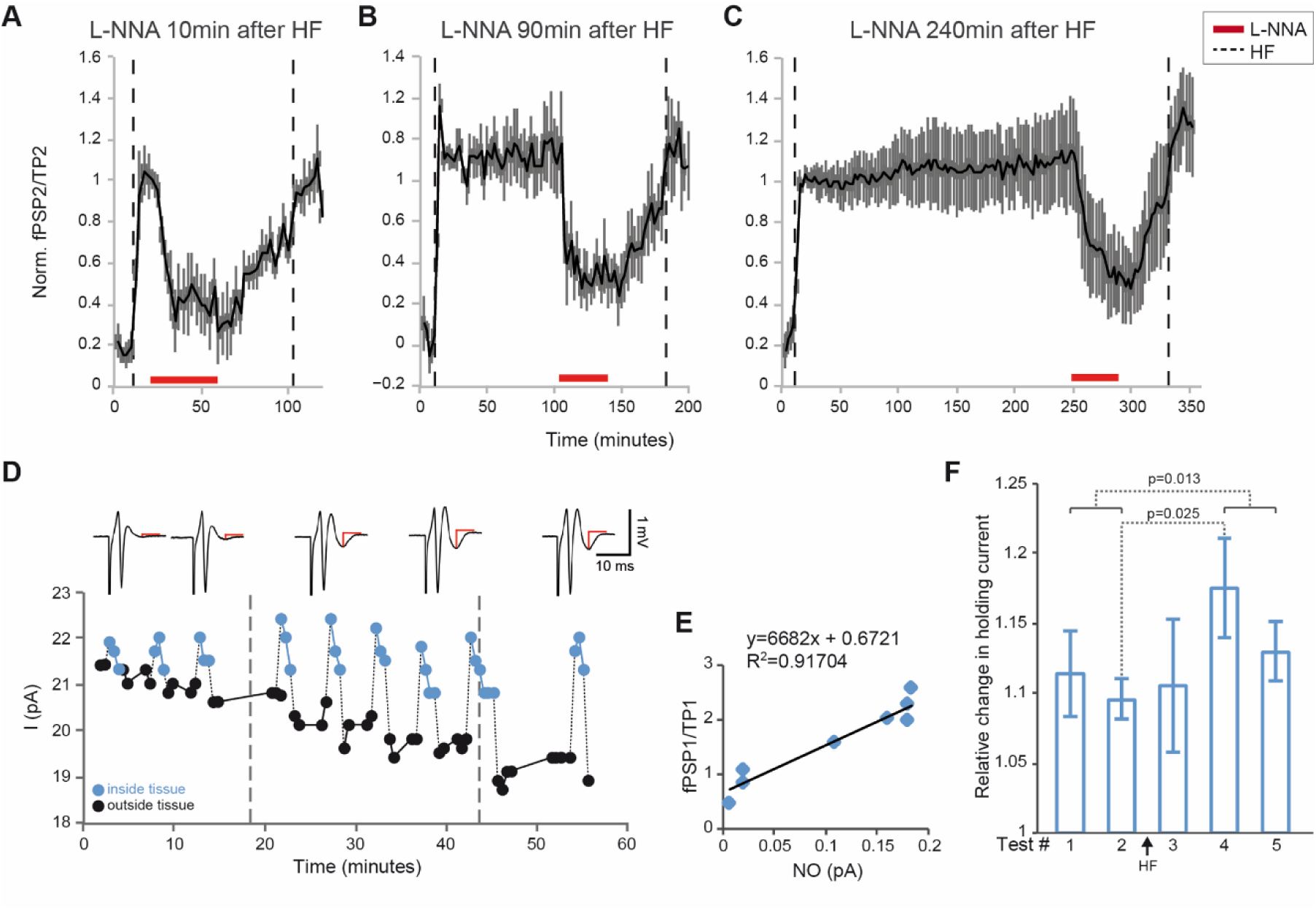
Persistent increase in NO concentration mediates the long-term presynaptic LTP *expression*. (A-C) L-NNA (10 mM) reversibly blocks LTP expression, irrespective of the time window after LTP induction: A, 10 min (n=4); B, 90 min (n=5); and C, 240 min (n=3). fPSP2/TP2 is normalized to LTP level obtained after the first HF stimulation. To detect residual LTP a second HF was delivered 40 min after starting L-NNA washout. (D-F) Amperometric measurement of NO concentration. (D) Current signals obtained by repeated insertions of the carbon fiber microelectrode (CFM) into the VL neuropil at the indicated times (blue dots). The corresponding fPSPs are shown in the upper insets (red lines mark fPSPs amplitude). The second HF stimulation was delivered when the electrode was inside the tissue. (E) A significant correlation between fPSPs amplitude and the amperometric NO signal as expressed by the difference between the currents measured inside and outside the neuropil (the same experiment shown in D). (F) Histogram depicting the average responses of five experiments like that shown in D. In each measurement (i.e., electrode insertion) the response magnitude was calculated by subtracting the average of two first current measurements following the withdrawal of the electrode from the average of first two measurements inside the tissue (blue dots in D). Each experiment was normalized to its maximal current response. Measurements were aligned with respect to the two tests before, and three tests after the HF stimulation (arrow). Data shown as mean ± SEM. Note a statistically significant delayed increase in NO signal following the HF stimulation as the first measurement after HF was not significantly different than the controls.

To directly test this possibility, we adopted an amperometric approach for measuring the extracellular concentration of NO (*Dias et al. 2016; Ferreira et al. 2005; Friedemann et al. 1996*) (*Figure 3D-F*). A NO-sensitive carbon fiber microelectrode (CFM, see Methods) was inserted for short periods into the VL neuropil. The experiment shown in Figure 3D started with three control insertions of the electrode. These insertions likely caused some gradual increase in the NO signals and a concomitant increase in fPSP (*Figure 3D insets*). After HF stimulation, the clamping current at the NO oxidation potential (750 mV) increased in parallel to the increase in the fPSP amplitude (*Figure 3D* and *insets*) – note the positive correlation between NO current and fPSP amplitudes (*Figure 3E*). The rough calibration of the CFM electrode, achieved by releasing NO from a known concentration of the NO donor SNAP, indicated an increase to approximately 0.5 µM. This concentration is about three orders of magnitude higher than the physiological range of NO concentrations (*Hall and Garthwaite, 2009*). As is typical for LTP saturation, neither the fPSP amplitude nor the NO signals were further increased by a second HF stimulation administered while the CFM electrode was inside the tissue (*Figure 3D*). Figure 3F shows the average relative reduction in the CFM electrode holding current measured inside the tissue (i.e., two first blue dots in *Figure 3D*) relative to that measured following withdrawal of the electrode (i.e., two first black dots in *Figure 3D*). The current signals increased substantially when the electrode was inserted into the isolated brain prior to the HF stimulation, likely due to some endogenous redox molecules. Yet, the average measurements after LTP induction show a significant elevation in the current of the second measurement after the HF stimulation relative to that measured before HF (t-test p = 0.025), indicating a possible delayed elevation in NO concentration. Although we demonstrated that NOS inhibitors blocked the amperometric response along with LTP expression, as expected, we also found that L-NNA (10 mM) blocked the CFM response to NO released from the NO donor SNAP. Consequently, we could not unequivocally confirm the involvement of NOS.

### NO-induced presynaptic facilitation requires unusually high NO concentrations

To directly test the involvement of NO in LTP expression and maintenance we used the NO donor SNAP. To our surprise, only mM ranges of SNAP (in the presence of 20 µM CuCl_2_ to facilitate SNAP decomposition; *Zhang et al. 2000*) produced an enormous synaptic facilitation (*Figure 4*). For example, Figure 4A, reveals a robust facilitation of fPSP/TP that was obtained with bath application of 2 mM SNAP (*Figure 4A* and *insets*). However, this facilitation was accompanied by a rapid decline in the TP amplitude that ended in a complete block of TP propagation, and consequently fPSP elimination likely due to the neurotoxic effect of the high NO concentration.

**Figure 4.**
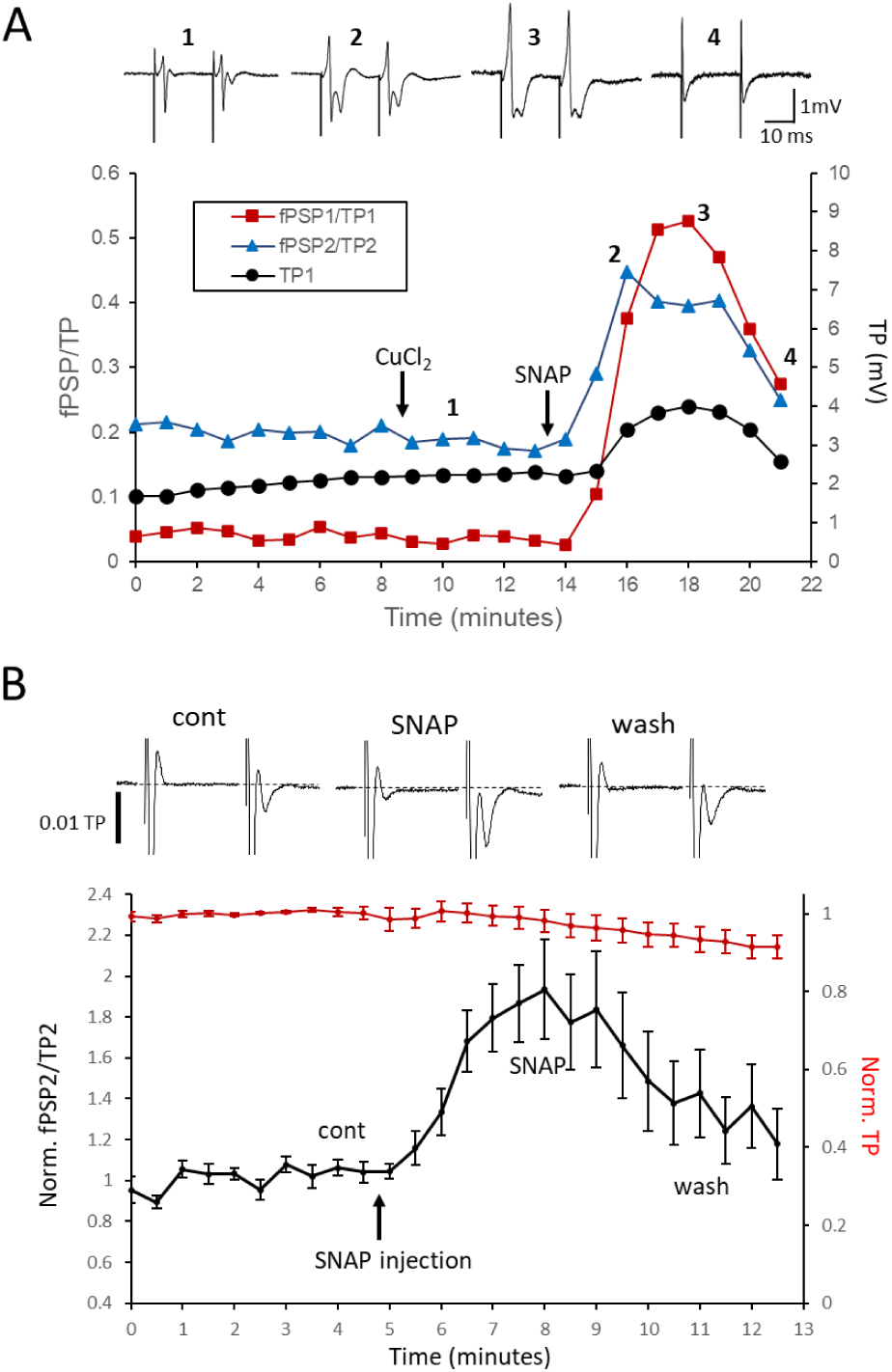
A considerably high range of NO concentrations is required for inducing short-term facilitation of transmitter release. (A) An example of the effects of exposing a slice preparation to millimolar concentrations of the NO donor SNAP (2 mM in the presence of 20µM CuCl_2_ to facilitate the release of NO) caused a dramatic transient facilitation of fPSP that was terminated due to the neurotoxic effect of NO on the tract action potential (insets correspond to the numbers on the graph). (B) Local pressure injections of sub microliters bolus of 10 mM SNAP, close to the recording site, induced short-term facilitation of fPSP/TP. The insets show the average fPSPs (the TPs are truncated) recorded in seven experiments (each experiment was first normalized to the amplitude of TP1 and then aligned at the negative peak of the TPs). The amplitude calibration bar (0.01 TP) represents the amplitude relative to TP=1. Note that fPSP1 is noticeable only during SNAP injection, while fPSP2 is transiently facilitated. In this procedure, higher NO concentration caused only short-term facilitation without the neurotoxic effects. Data shown as mean ± SEM.

Assuming that neurotoxic levels of NO are in the micromolar range (*Hall and Attwell, 2008; Hall and Garthwaite, 2009; Li et al., 2006*), it suggests that a local concentration of NO around µM is effective in inducing facilitation of transmitter release at the synaptic connections, corroborating the amperometric results. Similar to HF-induced LTP, SNAP-induced fPSP facilitation is also likely to involve an increased probability of transmitter release. This is indicated by the much larger relative increase in fPSP1/TP1 compared to fPSP2/TP2 (∼16-fold vs. ∼2.6-fold, respectively; *Figure 4A*).

The experiments illustrated in Figure 4B show that a brief pressure microinjection of a sub-microliter bolus of 10 mM SNAP resulted in short-term facilitation, which recovered after several minutes without causing any detrimental effects. Repeated injections produced additional transient facilitation responses. Importantly, these SNAP injections did not lead to a sustained fPSP potentiation, suggesting that NO alone is insufficient to induce LTP.

### The canonical NO-dependent c-GMP cascade does not appear to be involved in VL LTP

The high concentrations of SNAP required for the induction of synaptic facilitation together with the levels of NO concentration measured amperometrically, and the high concentrations of NOS inhibitors that were effective, suggest that the NO effects are mediated by much higher NO concentrations than the 100 pM to 5 nM effective in activation of the canonical NO-dependent soluble-guanylyl-cyclase (sGC) cGMP cascades (*Hall and Garthwaite 2009; Hardingham et al. 2013; Garthwaite 2016; Garthwaite 2019; Kleppisch and Feil 2009*). The recent results reported above may explain why our previous attempts to test the possible involvement of cGMP cascade have failed (*Turchetti-Maia et al. 2018*). Indeed, the NO scavenger (PTIO 1mM), shown to be effective in cephalopods chromatophore system (*Mattiello et al. 2010*), did not affect the potentiated fPSPs. This is likely because this NO scavenger is ineffective in significantly reducing the high NO concentration in the VL. Furthermore, attempts to assess the involvement of cGMP cascade in the VL LTP using a battery of pharmacological tools (listed in Methods) failed to intervene with either LTP expression or with its maintenance (*Turchetti-Maia et al. 2018*). Importantly these drugs were used successfully by Tu and Budelmann (1999; 2000 a,b) to demonstrate the involvement of NO-dependent cGMP cascade in the regulation of the cephalopods statocysts activity.

### LTP maintenance does not require *de novo* protein synthesis

Numerous studies indicate that the universal mechanism for maintaining the long-term phases of LTP involves *de novo* protein synthesis (*Kandel et al. 2014*) including serotonin-induced long-term facilitation in the mollusk Aplysia (*Montarolo et al. 1986*). In contrast we found that neither LTP induction nor its very long maintenance appears to involve protein synthesis. The administration of 20 µM anisomycin for 2.5 h, starting 30 min before the induction of LTP, blocked neither induction nor maintenance of LTP for at least 10 h after LTP induction (*Figure 5A*). We noticed that anisomycin may have side-effects which on one hand facilitated the induction of LTP even at the slow rate of the test stimulation (*Figure 5A,B*) and a slow untypical reduction in fPSP/TP ratio that was recovered following anisomycin washout (*Figure 5A,B*). This protein synthesis-independent LTP expression was blocked by L-NNA 10 mM, indicating that the mechanism for persistent activation of NOS was not affected by anisomycin (*Figure 5B*). Importantly, exposing *in vitro* slices to anisomycin 20 µM did suppress protein synthesis, including the intense protein synthesis in the VL (*Figure 5C*). The blocking effect of anisomycin 20 µM was replicated in five additional experiments.

**Figure 5.**
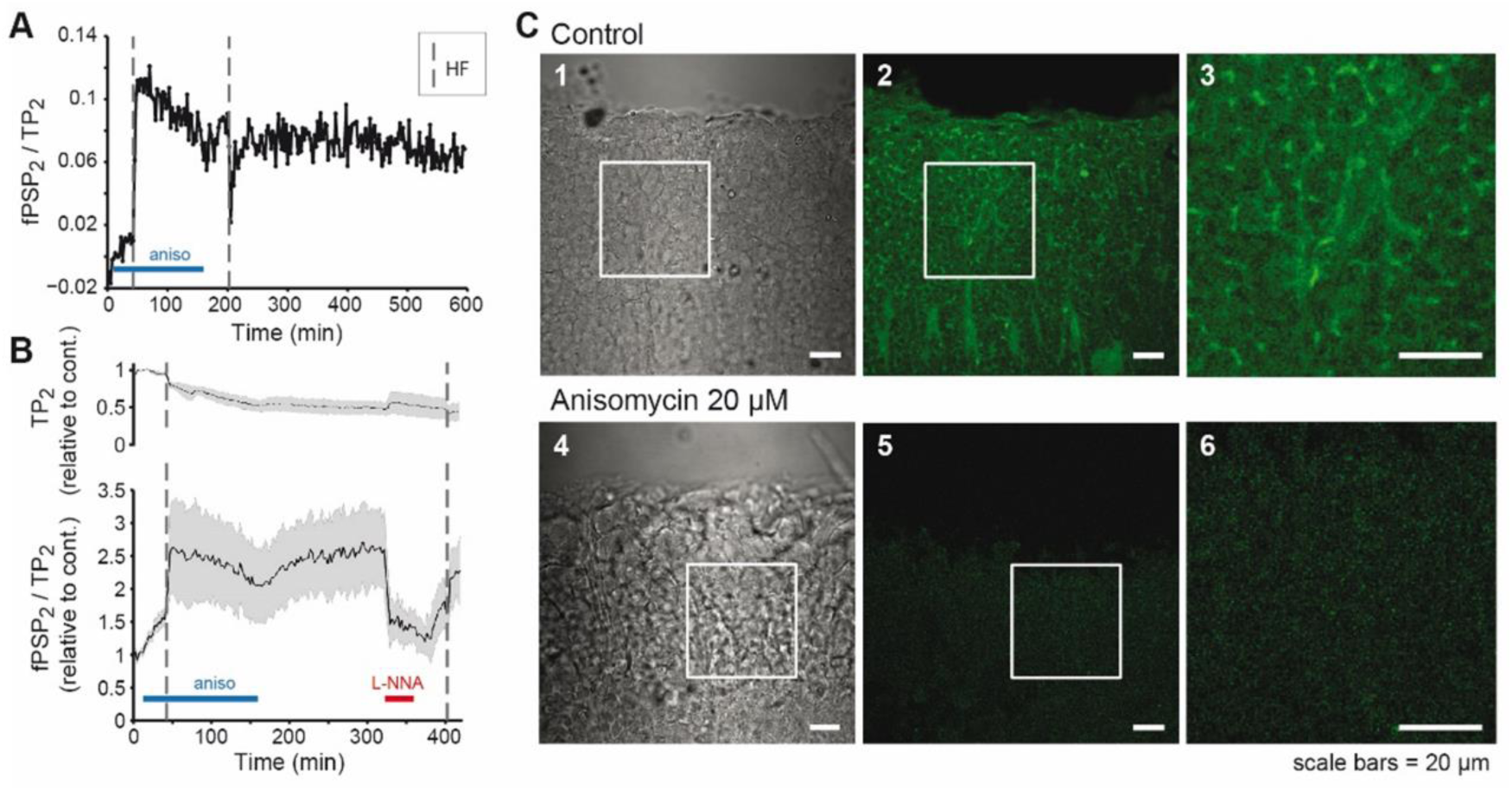
Long-term LTP *maintenance* is independent of *de novo* protein synthesis. (A-B) Testing the effect of exposure to anisomycin (20 μM, blue bars) on VL LTP induction and maintenance. (A) Example of a long duration experiment (10 h). A second HF stimulation, at 200 min, shows an intact LTP maintenance. (B) Population average of four experiments (mean ± SEM) showing the reversible effects of anisomycin on fPSPs/TP (lower graph) and TP amplitudes (upper graph). Note that L-NNA (10 mM, red bar) blocked the anisomycin-resistant LTP expression. (C) Confocal microscopy images of *de novo* protein synthesis using the FUNCAT assay (see Methods). C1 and C4: transmission bright-field images of the cell body cortex area. C2 and C3 are fluorescent images showing intense fluorescent signals, reporting protein synthesis, in control. C5 and C6 show that this signal is largely suppressed by anisomycin. The white squares in C1, C2 and C4, C5 mark the areas of magnification displayed in C3 and C6 respectively

### The NOS inhibitor L-NNA discriminates two types of LTP maintenance mechanisms

Figures 6A and B show the analysis of the kinetics of L-NNA-induced inhibition of LTP expression. For this *post-hoc* analysis, 11 of the 12 experiments shown in Figure 3A-C were aligned at 10 min before the onset of 42.5 min exposure to L-NNA (10 mM). We noticed dichotomic differences between experiments that showed either an abrupt potentiation induced by a second HF stimulation delivered during the drug washout (*Figure 6A, B*; blue curve, n=5, referred to as ’fast’) or those in which the second HF stimulation did not affect the rate of the monotonic recovery from inhibition (*Figure 6A, B*; red curve, n=6, referred to as ’slow’). The average potentiation induced in the ‘fast’ group was 2.54 ± 0.36 SE (n=5) (the average fPSP2/TP2 during the 10 min following the HF stimulation divided by the average in the 10 min before HF), while in the ‘slow’ group the average potentiation was 1.16 ± 0.07 (n=6). The obtained difference between the means was highly significant (t-Test; P = 0.015). Importantly, these two groups did not differ in their tract potential (TP) amplitudes (1.55 mV ± 0.25 SE vs. 1.63 mV ± 0.34 SE; t-test P= 0.853), thus suggesting that the two groups differ in their synaptic properties.

**Figure 6.**
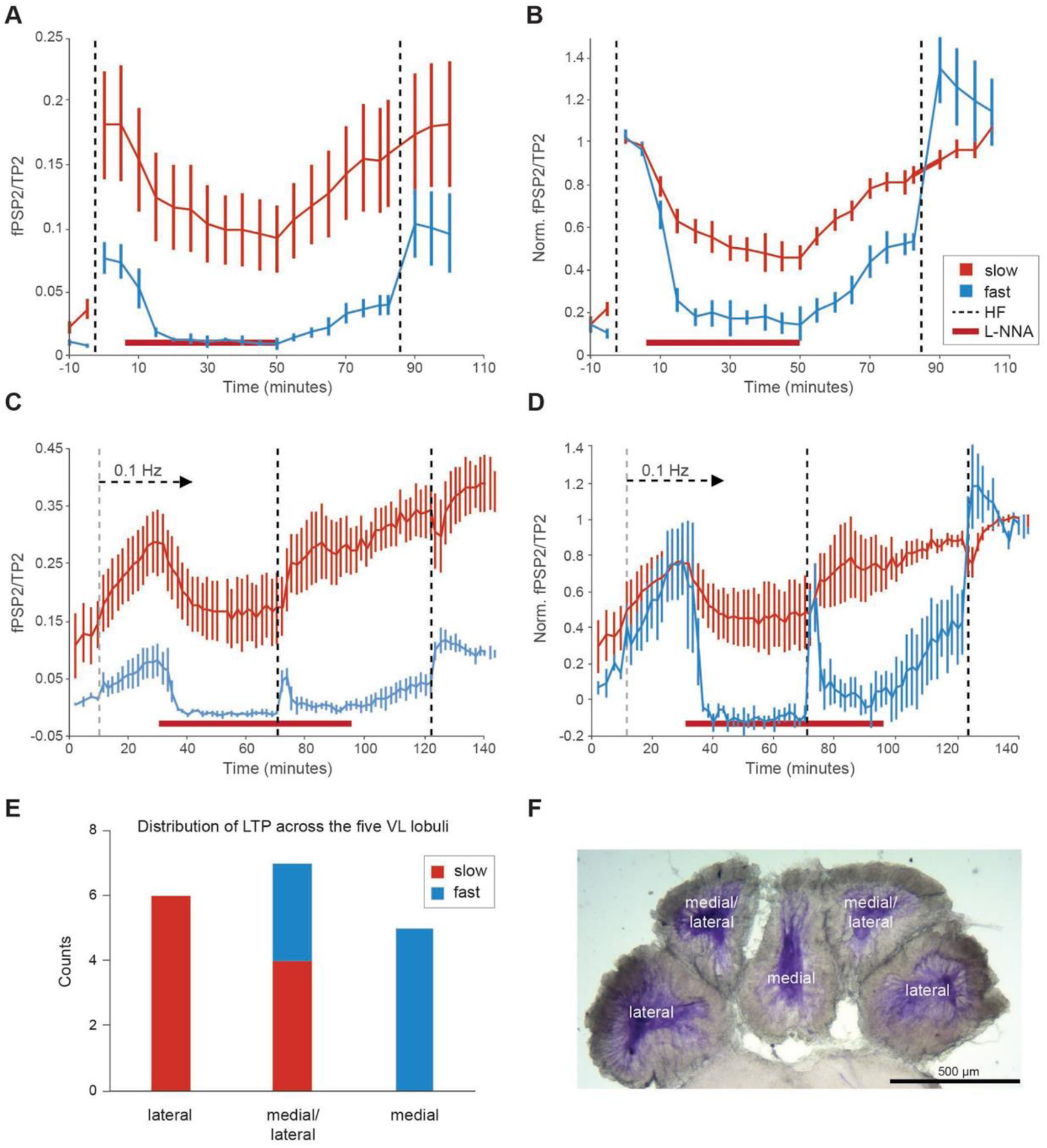
NO is involved in both LTP *expression* and *maintenance*. (A) Eleven experiments shown in Figure 3A-C in which 10 mM L-NNA was administered 10, 90 and 240 min after LTP induction were aligned at 10 min before the administration of the drug. Retrospectively this enabled segregating the inhibition kinetics into two groups (see text). (A) A group of large normalized fPSP (fPSP2/TP2) and a ‘slow’ inhibition time course (n=6; red curves), and a group of small fPSP and a ’fast‘ inhibition time course (n=5; blue trace). (B) The experiments shown in A were normalized to the level of LTP before L-NNA application. These depiction emphasizes the differences between the ‘slow’ and ‘fast’ inhibition kinetics and their response to a second HF which only in the ‘fast’ group induced abrupt potentiation. (C) Experiments in which the rate of the test pulses was increased from 0.033 Hz to 0.1 Hz (first dashed line). This rate of stimulation caused a slow development of LTP (SDP, slowly developed potentiation). Like in A, this set of experiments could be segregated into a large fPSP2/TP2, with slower time-course of inhibition (red; n=4) and small fPSP2/TP2 (blue; n=3) with a fast inhibition. HF stimulations were delivered during the steady state level of inhibition and at the recovery from the inhibition. (D) The same experiments as in C normalized to the level of LTP obtained after the second HF stimulation in order to emphasize the difference in inhibition and recovery kinetics. (E-F) Distribution of the two types of LTP across the five VL lobuli (the 18 experiments shown in A-D). (E) Histogram showing that ‘fast’ inhibition is prevalent in the medial lobule while ‘slow’’ inhibition is prominent in the two lateral lobuli. In the medial/lateral lobuli, both types exist.

Figure 6A,B show that the two groups are different with respect to three additional characteristics. First, the normalized fPSPs (fPSP2/TP2) of the ‘slow’ group (red curves), are much larger than those of the ‘fast’ group (3.5-fold in control and 2.3-fold following LTP induction) (*Figure 6A*). Second, in the ‘fast’ group, 10 mM L-NNA almost completely blocked LTP expression (97 ± 5.1%), bringing the fPSP amplitude back to the pre-LTP level (*Figure 6A,B*). In contrast, in the ‘slow’ group, the level of inhibition reached 72.1 ± 7.3%, which was significantly lower than the level of inhibition obtained in the fast group (two tail t-test equal variance P=0.024, with no difference in variance; f-test=0.403). These results indicate that in the ‘slow’ group, L-NNA reversibly inhibits NO-dependent LTP expression, while in the ‘fast’ group, L-NNA obliterates LTP maintenance, possibly by shifting NOS back to its inactivated state and therefore the second HF stimulation could activate a second round of HF induced LTP (*Figure 6A,B*). Note the gradual potentiation that appeared in the ‘fast’ group following L-NNA washout, is likely due to some recovery of expression and also to activity-dependent ‘slowly developed potentiation’ (SDP) induced by the test stimulations themselves (see below).

The third distinctive difference between the two groups is the large discrepancy between the rates of inhibition onset. Remarkably, LTP expression shutdown in the fast group is much faster than in the slow group (*Figure 6B*). The estimated exponential decay time-constant in the ‘fast’ group was more than four times faster (4.44 min; R^2^ = 0.985, Person correlation P = 0.008) than that of the ‘slow’ group (19.7 min; R^2^ = 0.939, Person P < 0.00001). While the slow time constant of the ‘slow’ group may reflect the rate of the drug perfusion, the fast and complete LTP shutdown in the ‘fast’ group suggests that this inhibition involves also an accelerated/regenerative deactivation of NOS. This strongly indicates the involvement of a positive feedback loop whereby NO reactivates NOS.

These two distinct groups of ‘slow’ and ‘fast’ inhibition kinetics were also clearly discerned in the *post-hoc* analysis of the experiments in which LTP was induced gradually by a moderate rate of stimulation (see Methods). These experiments are shown in Figure 6 C,D. Increasing the rate of the paired-pulse stimulation from 1/30 s to 1/10 s led to a slowly-developing-potentiation (SDP). Previously we suggested that this SDP shares the same mechanisms with that of HF-induced LTP because they occlude each other (*Shomrat et al. 2010*). The current work supports this conclusion as NOS inhibitors have blocking effects on SDP similar to that on HF induced-LTP. Here, also the inhibition kinetics by L-NNA (10 mM) could be segregated into the same two distinct groups as in the HF-induced LTP (compare *Figure 6 C,D* with *A,B*). In addition, three out of the seven SDP experiments had an exceptionally fast inhibition onset (blue curve) that fits an exponential time-constant of only 2.53 min (R² = 0.9491, Pearson P=0.0084). Consistent with the ‘fast’ group in HF-induced LTP, the ‘fast’ group in the SDP experiments also showed a complete blockage of SDP expression (*Figure 6 C,D*, blue). In this group, SDP maintenance was also blocked, as a second phase of SDP was initiated following L-NNA washout (around 100 min; *Figure 7C,D*) and HF-stimulation (given at 120 min) induced an abrupt potentiation. Note that in the ‘fast’ group HF stimulation given before drug washout caused a robust short-term potentiation that was terminated rapidly likely due to the fast inhibition kinetic of L-NNA. The slow SDP experiments *(Figure 6 C,D*, red) fit well with the ‘slow’ group in HF-induced LTP, exhibiting a slower inhibition time constant of 8.06 min (R² = 0.957; Pearson P = 0.005). In this ‘slow’ group, the L-NNA blocked only the SDP expression, as HF stimulation caused an abrupt and stable potentiation that was fully expressed after L-NNA washout, suggesting that HF-induced LTP can occur in the presence of NOS inhibitors (*Figure 2A*). Indeed, a second HF-stimulation (at 120 min) did not induce further potentiation, indicating that saturated LTP was induced by the first HF stimulation given during maximum inhibition (at 70 min). Like the HF-induced LTP experiments, shown in Figure 6A, here also a highly significant correlation exists between the ‘slow’ and ‘fast’ inhibition kinetics and the large and small fPSP2/TP2 amplitudes, respectively (*Figure 6C*). Overall, this dichotomic and large difference in the fPSP amplitude suggests that the ‘slow’ and ‘fast’ groups represent two different classes of SFL-to-AMs synaptic connections that also differ in their NO-dependent plasticity mechanism.

**Figure 7.**
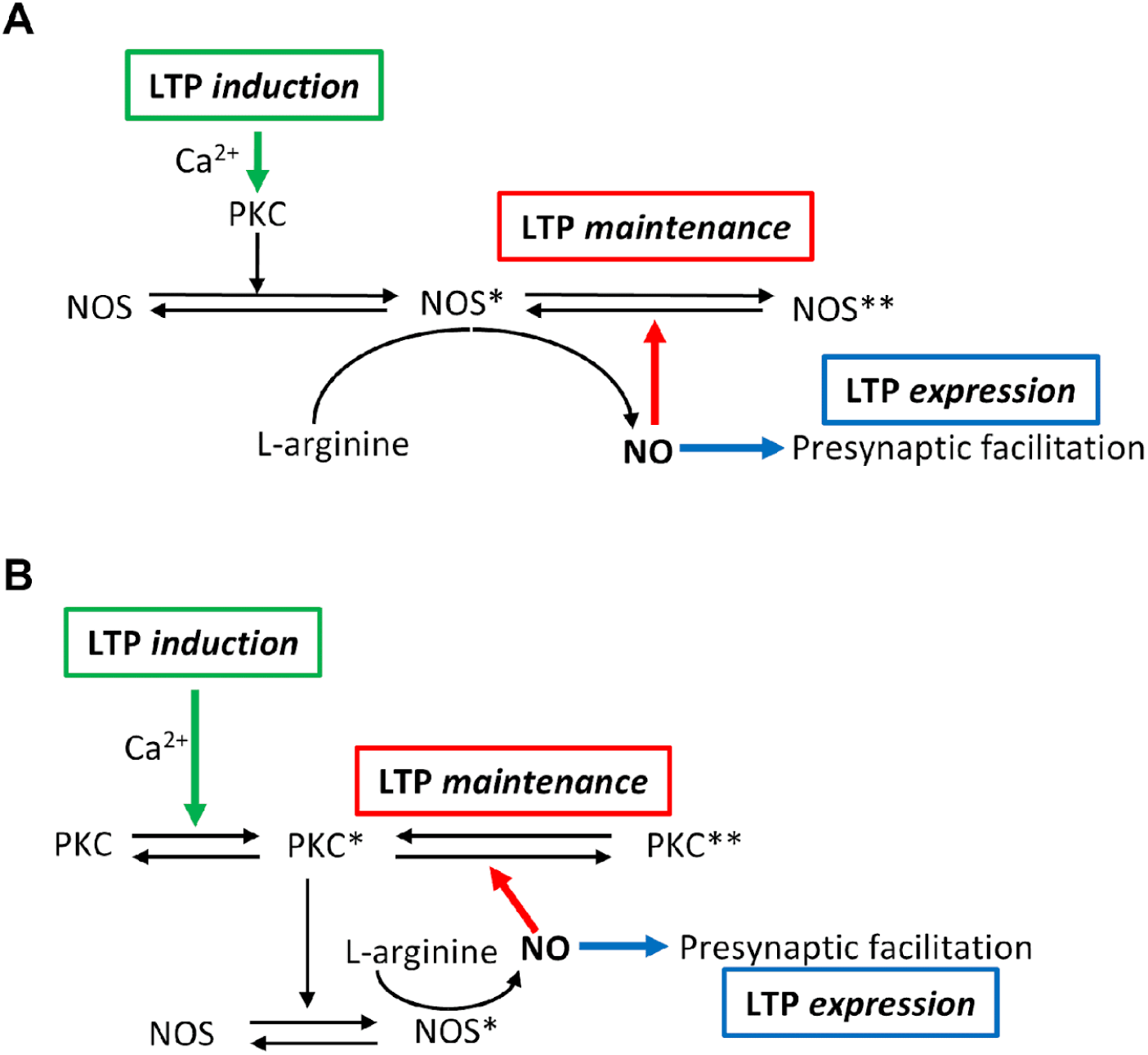
Two possible “molecular switch” models for the NO-dependent long-term *maintenance* and presynaptic *expression* of LTP in the octopus VL. For more details, see text.

Figures 6E,F show a cross section of the VL (taken from *Figure 1F*) labeled for NADPH-diaphorase which shows one medial, two medial-lateral, and two lateral lobuli. The histogram shows that the synaptic connection in the medial lobule represents only synapses with the ‘fast’ properties, while the lateral demonstrates the ‘slow’ properties. In the lateral/medial, both ‘fast’ and ‘slow’ properties could be documented. It should be pointed out that this classification is based on the slice number (typically 5-6 sagittal slices per VL) and therefore the correspondence with specific lobuli is only an approximation.

## Discussion

In this paper, we present a series of experiments revealing an unprecedented molecular mechanism that has evolved in the octopus brain to mediate activity-dependent long-term synaptic plasticity. Remarkably, this molecular mechanism, which is likely based on an adaptation of the molluscan nitrergic neuromodulation system, provides the octopus’ VL with cellular processes that, as we explain below, may support associative Hebbian plasticity, similar, in principle, to those provided by the NMDA-receptors in the mammalian hippocampus (*Malenka 2003; Malenka and Nicoll 1999*).

The effects of NOS inhibitors on LTP *expression* and *maintenance* (*Figures 2,3* and *6*) led us to propose a novel NO-dependent “molecular memory switch” mechanism. Figure 7A and B depict two possible molecular switch models. In both, activity elevates intracellular Ca^2+^ concentration which triggers LTP *induction* (green arrow) that involves Ca^2+^-dependent NOS activation in the postsynaptic cell (e.g., the SAMs). The generated NO then diffuses in a retrograde manner to the presynaptic side (blue arrow) of the synapse (SFL presynaptic boutons) where it induces facilitation of transmitter release (i.e. LTP *expression*). In parallel (red arrow), at least in the group of ‘fast’ synapses, NO also mediates *maintenance* through a positive feedback loop whereby NO dynamically “locks” NOS in active states, preventing NOS deactivation even when Ca^2+^ returns to its resting concentration (*Figure 7A*). The model shown in Figure 7B proposes that LTP induction is mediated indirectly through Ca^2+^-dependent PKC activation which in turn activates NOS. In model B, persistent activation of NOS is mediated by NO-dependent PKC activation. Indeed, we found that PKC activator (phorbol ester, PDBu 5 µM) induces long-term potentiation which, like LTP expression, can be blocked by NOS inhibitors (*Turchetti-Maia et al. 2016*).

To summarize, in the synapses with the ‘slow’ inhibition kinetics the competitive NOS inhibition reversibly blocks NO production thus diminishing LTP *expression*. In the synapses with the ‘fast’ and complete blocking kinetics the synergy between the reduction in NO concentration and the concomitant block of the positive feedback loop leads to the regenerative NOS deactivation. This reinstates NOS to its inactivated state and enables a new round of LTP following the removal of the inhibitor.

As the reversible inhibition of *expression,* and the irreversible block of *maintenance,* are correlated with strong and weak synapses, respectively (*Figure 6 A,C*), it is yet to be explored whether the two blocking modes are based on a single enzymatic cascade, which works differently depending on the strength of the postsynaptic response.

### NO mediates the presynaptic *expression* of LTP via a retrograde messenger pathway

We found that NOS activity is strongly expressed in the primary neurites of the majority of amacrine interneurons (AMs) (*Figure 1E* and see *Stern-Mentch et al. 2022*). In this wide range histolabeling study, Stern-Mentch et al. (2022) found that the primary neurites of substantial proportion of the AMs are labeled for choline acetyltransferase of the common type (cChAT) (*Casini et al. 2012*). These findings suggest that most cholinergic AMs also express NOS activity. Based on their morphological characteristics and their abundance, we conclude that these groups of NOS expressing cholinergic AMs belong to the group of simple amacrines (SAMs) that was characterized recently by Bidel and Meirovitch et al. (2023). It was discovered that 89.3% of the total of 25 million cell bodies in the VL cortex are SAMs because they possess un-bifurcating primary neurites that run into the neuropil (*Bidel and Meirovitch et al. 2023*).

The presence of NOS in the SAMs suggests that a sufficiently strong SFL-evoked postsynaptic response, such as during HF stimulation, activates NOS persistently in the SAM, leading to the continuous generation of NO (*Figure 7*). Then NO diffuses retrogradely to the presynaptic side of the activated synapse (the SFL axonal boutons), where it facilitates transmitter (glutamate) release. Interestingly, Bidel and Meirovitch et al. (2023) found that the SAMs receive only a single SFL input via a large presynaptic axonal bouton that synapses onto a palm-like postsynaptic structure on the SAM neurite. This structure likely increases the area of close apposition between the post- and presynaptic membranes, facilitating and restricting the retrograde diffusion of NO from the post-to the presynaptic cells ensuring ‘*synaptic specificity’*, a crucial property of associative networks.

It is tempting to speculate that selecting a NO-dependent process that works via volume transmission at very high NO concentration provides a sharp spatial gradient of effective NO concentrations. Such a virtue is conceivably important for ensuring *synaptic specificity* and *associativity* when plasticity is mediated by a retrograde messenger. An NO-dependent process that modifies proteins directly, for example through s-nitrosylation, may explain our findings. First, because we found no evidence for the involvement of the NO-dependent guanylyl cyclase cGMP cascade. And second, we found that the effective NO concentration in the VL is in the range which was reported to be required for direct protein s-nitrosylation modifications (*Hardingham et al. 2013; Hall and Garthwaite 2009*).

Ours and other literature investigations (e.g., *Padamsey and Emptage 2014; Susswein et al. 2004; Hardingham et al. 2013*) on the search for the involvement of NO as a retrograde messenger in presynaptic LTP are inconclusive. Perhaps the best evidence for such a mechanism was found in cultured hippocampal neurons (*Arancio et al. 1996; Arancio et al. 1995; O’Dell et al. 1991*). It is interesting to note that while the SFL-to-SAM synapses share morphological and cellular properties with the mossy fiber-to-CA3 pyramidal cell synapses (i.e., NMDAR-independent and presynaptic LTP expression), no experimental results support the involvement of NO as a retrograde messenger at this hippocampal synaptic connection. These findings correspond well with this synapse being the textbook example of non-associative LTP, and indeed, it was reported to be mediated by Ca^2+^-dependent PKA activation in the large mossy fibers’ presynaptic boutons (*Nicoll and Schmitz 2005; Vandael and Jonas 2024; Caya-Bissonnette and Béïque 2024*). Evidence for NO involvement in the heterosynaptic spread of presynaptically expressed LTP was suggested to occur at the synapses between the parallel fibers and Purkinje cells in the cerebellum (*Jacoby et al. 2001*).

### Can the molecular switch provide the mechanism for activity-dependent postsynaptically-dependent (Hebbian) associativity?

While LTP at a single SFL input into each SAMs can provide the essential learning rule of ‘*input (synaptic) specificity’*, it is harder to understand how plasticity at a one-to-one innervation pattern can provide the other essential rule of ‘*synaptic associativity*’ (*Malenka and Nicoll 1999; Malenka 2003*). This is especially intriguing because Shomrat et al. (2011) provided evidence that in the VL of *Octopus vulgaris*, LTP takes place at the “fan-out” SFL-to-AMs input layer and not at the “fan-in” AMs-to-LN output layer (see *Figure 1C*). Synaptic *associativity* requires a mechanism whereby only the synapses that are activated together will associatively facilitate each other’s likelihood to undergo LTP. Because we found that NO by itself induces only short-term facilitation (*Figure 4*), NOS will be “switched ON” into a persistent activation state only in the SAMs that are postsynaptically activated and NO concentration reaches some threshold (see molecular schemes in *Figure 7*). Given that the SAMs neurites intensely express NOS activity (*Figure 1D-F*) and that many SAM neurites are closely packed and fasciculate in a columnar organization (see *Bidel and Meirovitch et al. 2023*), it is therefore feasible that SAMs that are activated together will affect each other’s NO concentration via extrasynaptic volume transmission, leading to an associative “switching ON” of NOS.

This scenario may represent the “holy grail” constellation providing the means of both postsynaptic-mediated (Hebbian) synaptic *associativity* and synaptic *specificity* via the retrogradely NO-mediated presynaptic LTP *expression*.

## Methods

### Animals and electrophysiological recordings

Mature *Octopus vulgaris* of both sexes were collected by local fishermen on the Israeli Mediterranean coast. The conditions under which the animals were kept, the slice preparation procedures and the extracellular local field potential (LFP) recording methods followed Shomrat et al. (2011) unless mentioned otherwise. In some experiments we used isolated brain preparations that included the intact supraesophageal brain mass (*Shomrat et al. 2008*). The fPSP of the synapses between SFL axons and AM cells was evoked by stimulating the SFL tract at 0.03 Hz. LTP was induced by high frequency (HF) stimulation (4 trains of 20 stimuli at 50 Hz, 10 s inter-train interval). We routinely used paired pulses (20 ms apart) as test stimulation to study the paired pulse facilitation (PPF) ratio between the amplitudes of first and second fPSPs, quantitatively estimated by “PPF-index” (see details in *Figure 1*—*Figure supplement 1*). Unless otherwise indicated, the results are displayed as normalized second response (fPSP2/TP2), given that in most experiments under control conditions the first fPSP (fPSP1) was often indistinguishable from noise.

### Histochemistry

For NADPH diaphorase histochemistry, immediately after the supraesophageal mass was removed from anesthetized animals, the brain tissue was fixed by immersion in 4% paraformaldehyde (PFA) and 0.25% glutaraldehyde in artificial seawater (ASW) pH 7.4 at 4 °C overnight, then vibratome sectioned in the sagittal or transverse planes (50-100 µm). Tissue sections were analyzed for NADPH diaphorase activity according to Hope et al. (*Hope and Vincent 1989* modified by *Moroz 2000*, and see also *Stern-Mentch et al. 2022*). Briefly, slices were incubated at room temperature (RT) in the dark with reaction solution (0.5 M Tris-HCl buffer solution, pH 7.6, 1 mM β-NADPH, 0.2 mM Nitro Blue Tetrazolium (NBT), 0.3 % Triton X-100) for 30 min, rinsed with Tris-HCl buffer, then transferred to 4% PFA in methanol for one hour at RT. Finally, slices were dehydrated in ethanol, cleaned in xylene, mounted on SuperFrost Plus slides (Menzel Glaser, Germany) and viewed with Olympus SZX16 stereo and Olympus BX43 microscopes. Specificity of NADPH diaphorase staining was tested in control experiments in which tissue sections were incubated in the reaction solution as described, except that β-NADPH or NBT was omitted.

To evaluate anisomycin suppression of *de novo* protein synthesis in the octopus brain, we adapted the method of fluorescent noncanonical amino acid tagging (FUNCAT) using L-azidohomoalanine (Thermo Fisher Scientific) described in detail for hippocampal slices (*tom Dieck et al. 2012*). The adaptations aimed to guarantee marine invertebrate physiological conditions prior to fixation (ASW instead of Ringer solution, pH, and incubation temperature). Octopus slices were obtained as for electrophysiological recordings. We used Alexa Fluor 488 azide (Thermo Fisher Scientific) as fluorophore-alkyne-tag. Control slices were incubated with L-azidohomoalanine in the absence of anisomycin. Confocal imaging of the treated slices was performed using the Nikon C1 confocal system mounted on a Nikon TE-2000 Eclipse microscope system with a Nikon plan-Apo 60X NA 1.4 objective. Images were collected and processed using EZ-C1 software (Nikon). The Alexa Fluor 488 azide was excited with the 488 nm laser and the emission was collected with a 515 ± 30 nm filter. Images were prepared using NIH ImageJ software (Bethesda, MD, USA).

### Amperometry

NO was measured directly in the VL using electrochemical methods with 30 µm polypropylene-insulated carbon fiber microelectrodes (CFMs) (ProCFE Dagan Corporation) coated by dipping the electrode tip five times into Nafion® 117 solution and drying the electrode for 10 min in 85 °C (*Dias et al. 2016; Friedemann et al. 1996*). This procedure was repeated ten times. The CFMs were then coated by electro-polymerization with o-phenylenediamine dihydrochloride (Friedemann et al. 1996) and cellulose (*Marinesco et al. 2003*). All these coatings were found to be essential for increasing the sensitivity and specificity of the CFM to NO in ASW. To test the electrodes, we measured the release of NO from a known concentration of NO donor (stock solution of SNAP prepared in 100 µM EDTA). 10 µM CuCl_2_ was added to the ASW to catalyze SNAP decomposition to NO and disulfide by-product (*Zhang et al. 2000*). This concentration of Cu^2+^ had no effect on the physiological responses. For testing the peak oxidation voltage of NO in ASW, a slowly rising (∼2.3 s) voltage ramp from a DC holding voltage of 0.5 V to 1.1 V was used as a command for clamping the CFM potential with a Chem-Clamp Voltmeter-Amperometer (Dagan Corporation). Adding 10 µM SNAP to ASW containing 10 µM CuCl_2_ caused a maximal positive current response of close to 750 mV. Hence, for the *in vitro* measurements of NO during physiological experiments the electrode potential was clamped at a constant 750 mV. Because the oxidation current declined gradually in the presence of constant NO concentration, we used a micrometric manipulator to insert the CFMs into the tissue (∼100 µm deep) every 5 min for only 90 s. Amperometric measurements were stored together with the recording of the LFP generated by stimulation of the SFL tract. To calculate population results (*Figure 3F*), we took the average of three measurements inside the tissue minus the holding current following withdrawal of the electrode. Each experiment was normalized to the maximal current response.

All the *in vitro* experiments utilized isolated brain preparations (see *Shomrat et al. 2008*) comprising the supraesophageal part of the central brain (*Figure 1*). At the end of each experiment the CFM sensitivity in ASW was estimated by adding 10 µM SNAP to the experimental bath containing 10 µM CuCl_2_.

### Drugs

Drugs were administered via the perfusion system, taking ∼1 min to reach the recording site. We used the NOS inhibitors L-NNA (*N*ω-Nitro-L-arginine, 10 mM) and L-NAME (*N*ω-Nitro-L-arginine methyl ester hydrochloride, 10 mM), and its inactive enantiomer D-NAME (*N*ω-Nitro-D-arginine methyl ester hydrochloride, 10 mM). We routinely used 10 mM of these drugs because we found that at lower than 5 mM the results were less consistent. We found that obtaining 10 mM L-NNA in ASW, a 30 min of continuous stirring was required. We also used the cGMP inhibitors ODQ (1H-[1,2,4] Oxadiazolo[4,3-a]quinoxalin-1-one; 50μM) and methylene blue (0.1 mM); the cGMP agonists dBcGMP (0.2mM), 8-Br-cGMP (0.1mM), 8-pCPT-cGMP (10-100μM); the cGMP phosphodiesterase inhibitor ZAP (Zaprinast; 0.1mM); the cyclic nucleotide phosphodiesterase inhibitor IBMX (3-Isobutyl-1-methylxanthine; 0.5-1mM); the NO donors SNAP (S-Nitroso-N-acetyl-DL-penicillamine; 1-100µM and up to 10mM), DETA (Diethylenetriamine NONOate; 0.4 mM), and DEA (Diethylamine NONOate; 0.1mM); the NO scavenger PTIO (2-Phenyl-4,4,5,5 tetramethylimidazoline-1-oxyl 3-oxide; 1 mM). Some drugs were dissolved in ASW and a fresh solution made before the experiments (L-NNA, L-NAME, D-NAME, methylene blue). Stock solution aliquots were prepared in DDW (dBcGMP), in DMSO 0.1-0.25% (ODQ, ZAP, IBMX), in ethanol 1% (PTIO), in NaOH 0.05 mM (DEA), or in 100 µM EDTA (SNAP), and were diluted in ASW before the experiments in a concentration in which the vehicles had no physiological effects. When required, pH was adjusted to 7.6 using NaOH or HCl. Slices treated with reversible drugs, which could be washed out, could be used a second time in a different recording location to guarantee a different and yet unstimulated local population of cells. All drugs and chemicals were purchased from Sigma-Aldrich, unless stated otherwise.

### Data analysis

Electrophysiological data were analyzed as described in Shomrat et al. 2011. Unless otherwise stated, we used averages of 5 trials for all the electrophysiological recordings (30 s interstimulation interval). In the experiments where the slices were exposed to the drug before LTP induction, we normalized the responses relative to the control baseline (first 7.5 min of the recording). Where we sought to quantify the degree of LTP blockage, LTP was induced prior to drug exposure and the responses were normalized relative to LTP (7.5 min recording after HF). Values are presented as mean ± SEM.

## Supporting information

Supplemental methods and analysis Fig

## Acknowledgments

We thank Hadas Erez and Micha Spira for helping with the confocal images. A.L.T.M. received postdoctoral fellowships from the Lady Davis, Golda Meir Foundations, and The Edmond and Lily Safra Center for Brain Science (ELSC). This research was carried out through an extended period of time and was supported by the Israel Science Foundation (1425/11; 1928/15; 2937/21) the US-Israel Binational Science Foundation (2011-466), the Human Frontier Science Program (HFSP) RGP0042/2019 and was aided by the Smith Family Laboratory at HUJI.

## Footnotes

Author Contributions: A.L.T.M., T.S. and B.H. designed the study; A.L.T.M. performed and analyzed the physiological experiments with contribution from T.S. and B.H.; T.S. and B.H. designed and performed the NO-donor experiments; N.S.M., T.S. and N.N. designed and performed the NADPH diaphorase experiments; A.L.T.M., N.S.M., F.B. and N.N. performed the protein synthesis fluorescent assay; B.H. and T.S. performed the amperometric experiments and analyzed the results; A.L.T.M. T.S and B.H. wrote the paper with comments from all the authors.

The authors declare no competing interests.

