## Supplemental methods and analysis Fig for "A novel nitric oxide (NO)-dependent ‘molecular switch’ mediates LTP in the *Octopus vulgaris* brain through persistent activation of nitric oxide synthase (NOS)"

### *Figure 2 —figure supplement 1*. The basic properties of activity-dependent LTP in slice preparation of the octopus VL.

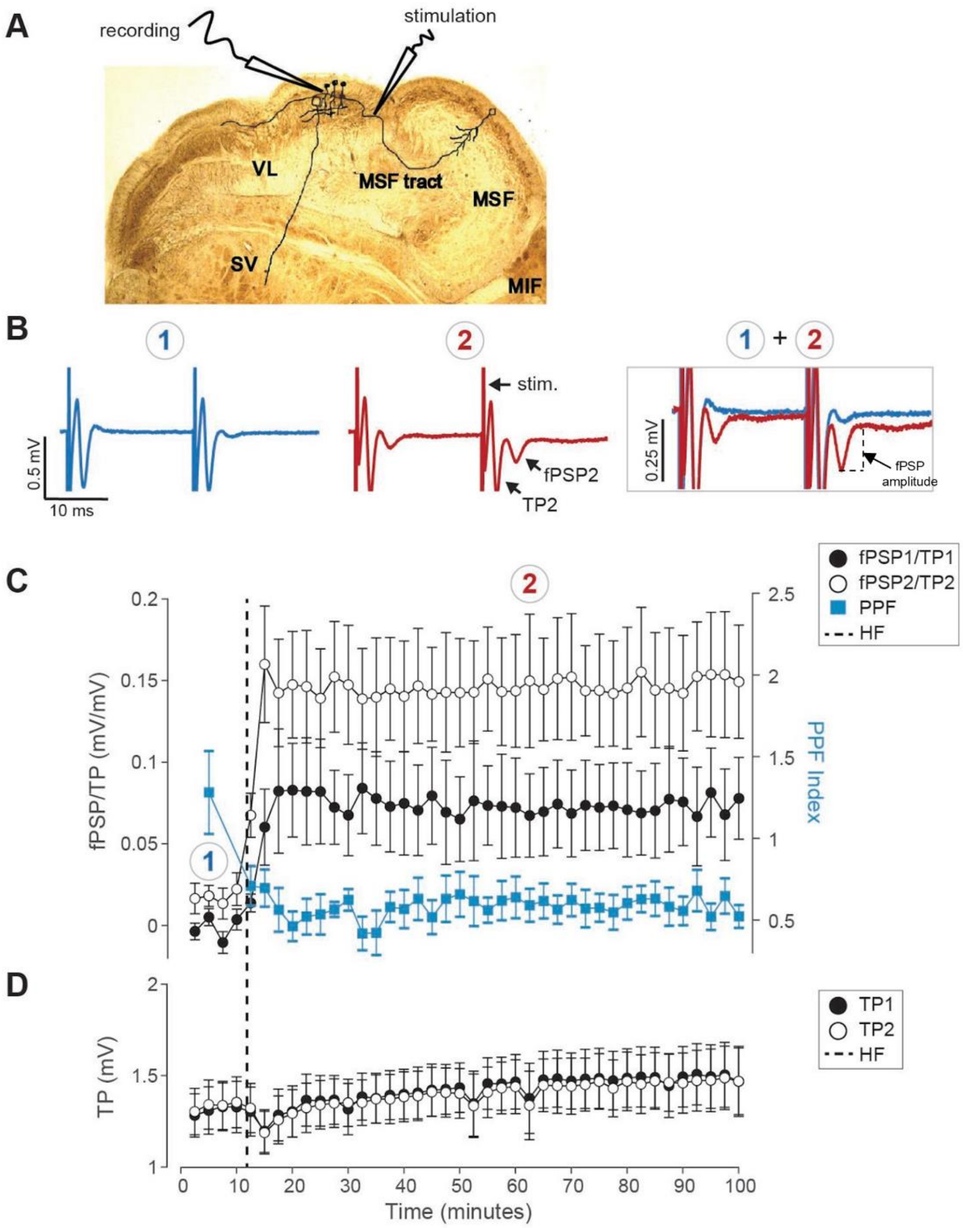


As previously described (*Hochner et al. 2003; Shomrat et al. 2011*), the stimulating and recording electrodes were placed on the SFL tract at a short distance (~ 0.5 mm) from each other (*Figure 2—figure supplement 1A*). The tract was stimulated with paired test pulses and the evoked tract potential (TP) and the generated synaptic field potential (fPSP; *Figure 2—figure supplement 1B*) were recorded. LTP was induced by high frequency stimulation (HF, four trains of 20 pulses at 50 Hz, 10 s inter-train interval). This HF stimulation led to a robust and long-term potentiation of the first and the second synaptic fields' potentials (fPSP1, fPSP2) (*Figure 2—figure supplement 1B,C*) with no significant change in the respective evoked tract potentials (TP1, TP2) (*Figure 2—figure supplement 1B,D*). Postsynaptic responses were measured as the fPSP amplitude (see dashed line *Figure 2—supplement figure 1B*). Likely due to the inexcitability of the AMs, the fPSP amplitude is linearly related to the TP amplitude (see *Hochner et al. 2003; Shomrat et al. 2011*). Therefore, to compensate for small changes in TP amplitude during the experiments, the fPSP was normalized relative to the TP amplitude (i.e., fPSP/TP) (*Figure 2—figure supplement 1C*).

The development of LTP is characterized by a change in paired-pulse facilitation (PPF), expressed here by PPF-index (PPF-index = (fPSP2/TP2 - fPSP1/TP1)/(fPSP2/TP2); *Figure 2—figure supplement 1C* blue squares and right vertical axis). The reduction in PPF-index indicates that the octopus VL LTP involves presynaptic increase in probability of transmitter release (*Manabe et al. 1993; Yang and Calakos 2013; Hochner et al. 2003*). While theoretically PPF-index allows quantification of PPF even when the fPSP1 is close to zero (i.e., PPF-index = 1), as often occurs, especially before LTP induction, fPSP1 is indistinguishable from the noisy background and therefore the PPF index is highly variable but yet not significantly different than PPF-index = 1.0 (1.288 ± 0.67 SEM; *Figure 2—figure supplement 1C*, blue squares). Following LTP induction, the relative increase in the amplitude of fPSP1 is the most prominent (see superimposed traces in *Figure 2—figure supplement 1B*) and the average PPF index is 0.584 ± 0.282 SEM, significantly lower than 1 (*Figure 2—figure supplement 1C*, right vertical axis).
